# Hormonal signaling cascades required for phototaxis switch in wandering *Leptinotarsa decemlineata* larvae

**DOI:** 10.1101/325233

**Authors:** Qing-Wei Meng, Qing-Yu Xu, Tao-Tao Zhu, Lin Jin, Kai-Yun Fu, Wen-Chao Guo, Guo-Qing Li

**Affiliations:** Education Ministry Key Laboratory of Integrated Management of Crop Diseases and Pests, College of Plant Protection, Nanjing Agricultural University, Nanjing 210095, China; Department of Plant Protection, Xinjiang Academy of Agricultural Sciences; Urumqi 830091, China; Xinjiang Laboratory of Special Environmental Microbiology; Institute of Microbiology, Xinjiang Academy of Agricultural Sciences; Urumqi 830091, China

## Abstract

Many animals exploit several niches sequentially during their life cycles, a fitness referred to as ontogenetic niche shift (ONS). To successfully accomplish ONS, transition between development stages is often coupled with changes in one or more primitive, instinctive behaviors. Yet, the underlining molecular mechanisms remain elusive. We show here that *Leptinotarsa decemlineata* larvae finish their ONS at the wandering stage by leaving the plant and pupating in soil. At middle wandering phase, larvae also switch their phototactic behavior, from photophilic at foraging period to photophobic. We find that enhancement of juvenile hormone (JH) signal delays the phototactic switch, and *vise verse*. Moreover, RNA interference (RNAi)-aided knockdown of *LdPTTH* (prothoracicotropic hormone gene) or *LdTorso* (PTTH receptor gene) impairs avoidance response to light, a phenotype nonrescuable by 20-hydroxyecdysone. Consequently, the RNAi beetles pupate at the soil surface or in shallow layer of soil, with most of them failing to construct pupation chambers. Furthermore, a combination of depletion of *LdPTTH/LdTorso* and disturbance of JH signal causes no additional effects on light avoidance response and pupation site. Finally, we establish that TrpA1 (transient receptor potential (TRP) cation channel) is necessary for light avoidance behavior, acting downstream of PTTH. We conclude that JH/PTTH cascade concomitantly regulates metamorphosis and the phototaxis switch, to drive ONS of the wandering beetles from plant into soil to start the immobile pupal stage.

**Author summary:** Many animals occupy distinct niches and utilize diverse resources at different development stages in order to meet stage-dependent requirements and overcome stage-specific limitations. This fitness is referred to as ontogenetic niche shift (ONS). During the preparation for ONS, animals often change one or more primitive, instinctive behaviors. Holometabolous insects, with four discrete developmental periods usually in different niches, are a suitable animal group to explore the molecular modes of these behavioral switches. Here we find that *Leptinotarsa decemlineata* larvae, an insect defoliator of potatoes, switch their phototactic behavior, from photophilic at feeding period to photophobic during the larval-pupal transition (wandering stage). This phototactic switch facilitates the wandering larvae to accomplish the ONS from potato plant to their pupation site below ground. We show that JH/PTTH cascade controls the phototaxis switch, through a step in photo transduction between the photoreceptor molecule and the transient receptor potential cation channel.

## Introduction

Movements to stage-dependent resources, i.e., ontogenetic niche shifts (hereafter ONS), occur in nearly 80% of animal taxa. The shifts enable animals to exploit several niches sequentially during their life cycles to meet stage-dependent nutritional requirements, to overcome stage-specific physiological limitations, and to reduce intraspecific competition between juveniles and adults. ONS is thus widely accepted as an evolutionary adaptation [1–4]. To successfully finish ONS, transition between development stages is often accompanied with changes in one or more primitive, instinctive behaviors, allowing inexperienced novices to obtain novel abilities to detect new environmental cues [5]. To date, however, the underlining mechanisms driving these behavior switches are still largely unexplored.

Insects are a suitable animal group to explore the molecular modes of these instinctive behavioral switches. Throughout developmental excursion, most insects (Holometabolans) undergo four discrete periods (complete metamorphosis), characterized by the presence of pupal stage between a feeding larva and a reproducing adult [6,7]. Sessile pupae are vulnerable to potentially harmful factors such as desiccation, predation, parasitism and pathogen infection. These latent mortal dangers drive a lot of Holometabolans shifting into less risky habitats for pupation [8–12]. For example, pupation in soil and other relatively inaccessible sites for predators and parasitoids (concealed placement) has been documented in almost all Holometabolan orders [8, 12–15].

Insect final instar larval stage is divided into two sub-stages, the foraging and the wandering phases. While foraging final instar larvae display generally similar behaviors like previous instar animals, a wandering larva typically undergoes an ONS by leaving the food source and moving to a proper pupation site [16]. Obviously, positive phototaxis directs most foraging insect herbivores to reach plant top for tender plant tissues such as shoots, young leaves, buds and flowers that are more nutritious and less defended [17], whereas negative phototaxis facilitates the wandering larvae to reach pupation refuge in the dark, such as soil [5]. Accordingly, it can be reasonably hypothesized that the change from foraging to wandering stages should be coupled with a switch for phototactic behavior from photophilic to photophobic in most herbivorous Holometabolans.

For a soil-pupated insect species, a wandering larva usually shows a sequence of three primary behavioral components before pupation: a) crawling to the ground and searching for a suitable location, b) mining into soil, and c) building a pupation chamber in soil [18–20]. Ecdysteroids (the major active component is 20-hydroxyecdysone, 20E), the products in a pair of prothoracic glands (PGs), activate the wandering behavior [18, 21–23].

Up to now, however, the molecular mechanism elicits the phototaxis switch in insect herbivores remains elusive. In *Drosophila melanogaster*, larval phototaxis and behavioral responses have been described [5, 24–27]. Unfortunately, photophobic is age-independent in the larvae [5, 24]. At the foraging stage, a larva feeds inside rotting fruits; it is strongly repelled by light and seeks for dark or less light-exposed surroundings [5, 25–27]. Two pairs of neurons called NP394 (each pair in a hemisphere of central brain) are required to maintain light avoidance in the foraging phase. Modulating activity of NP394 neurons affects larval light preference [28]. The NP394 neurons turn out to be prothoracicotropic hormone (PTTH)-expressing cells [5]. These two pairs of PTTH-producing neurons release PTTH to concomitantly promote steroidogenesis and light avoidance during wandering stage of the final instar larvae [5]. On one hand, PTTH, through its receptor Torso, activates a canonical mitogen activated protein kinase (MAPK) pathway to trigger ecdysteroidogenesis by PGs to regulate metamorphosis [29–31]. On the other hand, PTTH/Torso complex acts on two light sensors, the Bolwig’s organ (a group of 12 photoreceptors in the larval eye) and the peripheral class IV dendritic arborization neurons, to reinforce light avoidance [5].

The young larvae of the Colorado potato beetle *Leptinotarsa decemlineata*, a notorious insect defoliator of potatoes, reveal a tendency to rest and feed on the upper surfaces of leaves during foraging stage [32]. At the late stage of the final (fourth) instar, conversely, the wandering larvae bury themselves into soil, where, after several days, they metamorphose into pupae [33, 34]. Moreover, by RNA interference (RNAi), we have identified major components of the hormonal network that regulates larval metamorphosis in *L. decemlineata* [33–41]. This offers a great opportunity to explore the molecular mode driving the phototaxis switch in an insect herbivore.

The first aim of the current study was to determine whether *Leptinotarsa* larvae changed their light preference from photophilic to photophobic during wandering stage. We then uncovered that JH/PTTH cascade concomitantly regulated metamorphosis and the phototaxis switch. Finally, we provided clear evidence that PTTH-induced light avoidance drove ONS from plant to pupation refuge in an insect herbivore.

## Results

### Phototaxis of *L. decemlineata* larvae

We first observed phototaxis of *Leptinotarsa* beetles on the natural potato field. The females unselectively deposited their egg masses on upper and lower leaf surfaces (for a total of 100 egg masses 48 and 52 respectively on upper and lower surfaces, P>0.05 for χ^2^ test) at the inner part of the potato canopy (Figs 1A, B). Aggregated hatchlings consumed foliage near the egg mass from which they hatched (Fig 1C). All the second-, third-and foraging fourth-instar larvae were found to feed and rest on the upper surfaces of the potato leaves (Figs 1D-F). These larvae molted under sunny lighting condition (Fig 1F). During the wandering larval stage from approximately 4.1 to 7.0 days after ecdysis to the fourth instar (Figs 1G-I), the animals left the potatoes and crawled to the ground, walked along the ground (4.5-5.5 days post ecdysis), dug into soil, constructed their pupation chambers and pupated therein (5.6-7.0 days) (Figs 1G-I).

**Fig 1.**
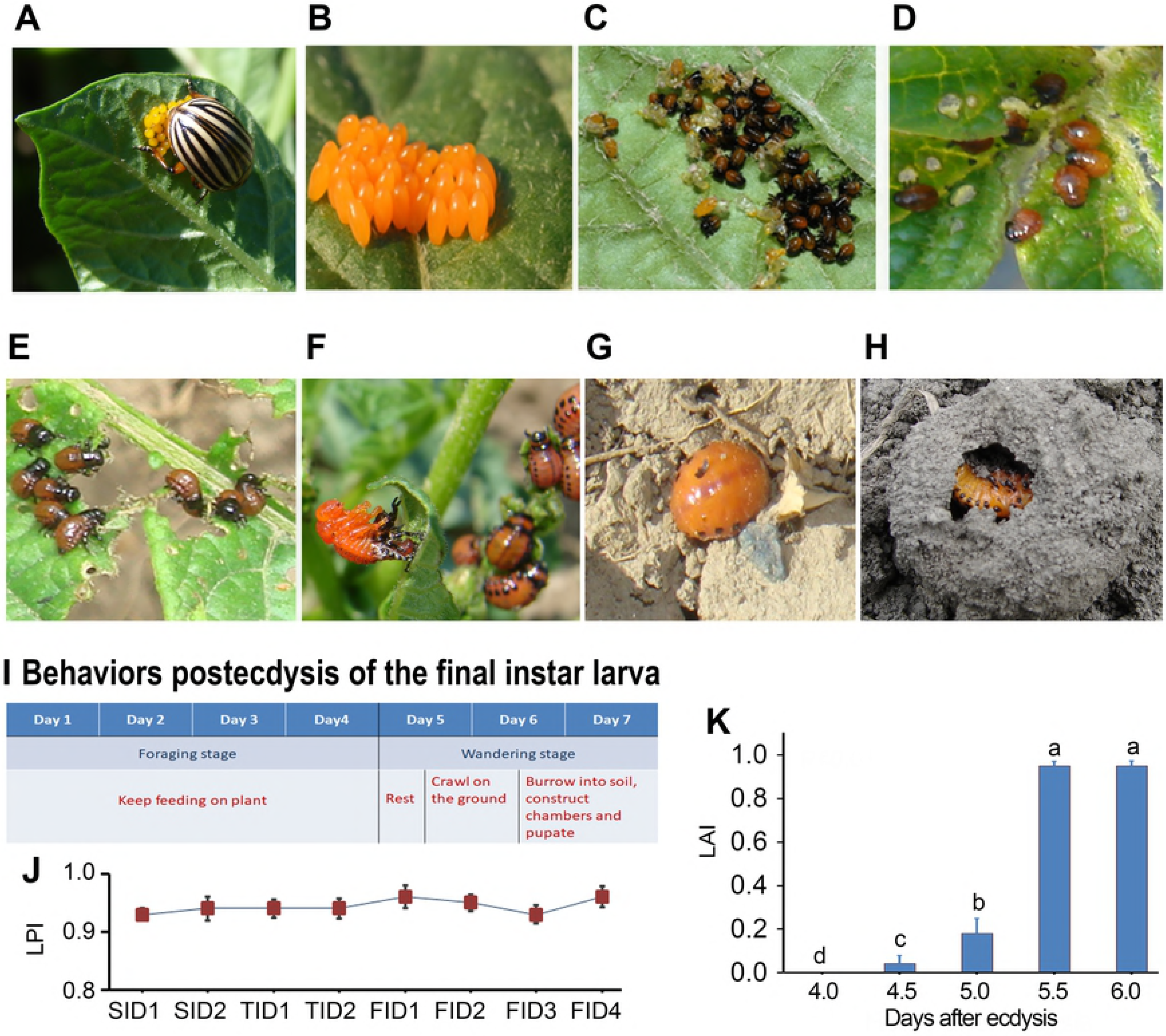
A phototaxis switch of the wandering larvae. (A-I) A total of 100 *Leptinotarsa* egg masses were selected randomly along a diagonal line. The location sites of egg masses (A and B), hatchlings (C), second-and third-instar larvae (D, E), fourth-instar larvae and a larva is molting to fourth instar (F), wandering larvae (G) and prepupae (H) were observed. The duration from ecdysis to pupation of the final instar larvae last about 7.0 days (I) (J, K) A two-choice test in laboratory. Light preference indexes (LPI) were assessed for second-instar (SI), third-instar (TI) and foraging fourth-instar (FI) larvae (D1-D4 indicates day 1 to day 4 after ecdysis) (J). Light avoidance indexes (LAI) were assessed at each testing time point from 4.0 to 6.0 days at an interval of a half day (different letter indicating significant differences among different time point at P<0.05) (K). (TIF)

When given a choice in the laboratory, second-instar, third-instar and foraging fourth-instar larvae prefer light-exposed over shaded areas (Fig 1J). For wandering *Leptinotarsa* larvae, however, light avoidance index significantly increased at 5 days post ecdysis, and reached nearly 1.0 at 6 days (Fig 1K). These basic findings demonstrate that an obvious phototactic switch occurs at the middle wandering period.

### Juvenile hormone signal inhibits the phototaxis switch

In *L. decemlineata*, transition from foraging to wandering phases is associated with drastic level fluctuations of three larval hormones: 20-hydroxyecdysone (20E), insulin-like peptide (ILP) and juvenile hormone (JH) [33, 34, 36, 40]. Are these hormone signaling cascades involved in the regulation of phototaxis switch?

In the final larval instar of *L. decemlineata*, a small 20E rise appears at 4 days after ecdysis [42]. Here, we found that dietary supplement with 20E to generate a premature 20E peak at the foraging stage (Fig 2A), or knockdown of an ecdysteroidogenesis gene [38] to remove this 20E rise (Fig 2B and S1A Fig), did not affect the phototaxis switch, neither did silencing of a 20E receptor gene *EcR* (Fig 2C and S1B Fig) or a 20E cascade gene *HR3* (Fig 2D and S1C Fig) to eliminate 20E signal.

**Fig 2.**
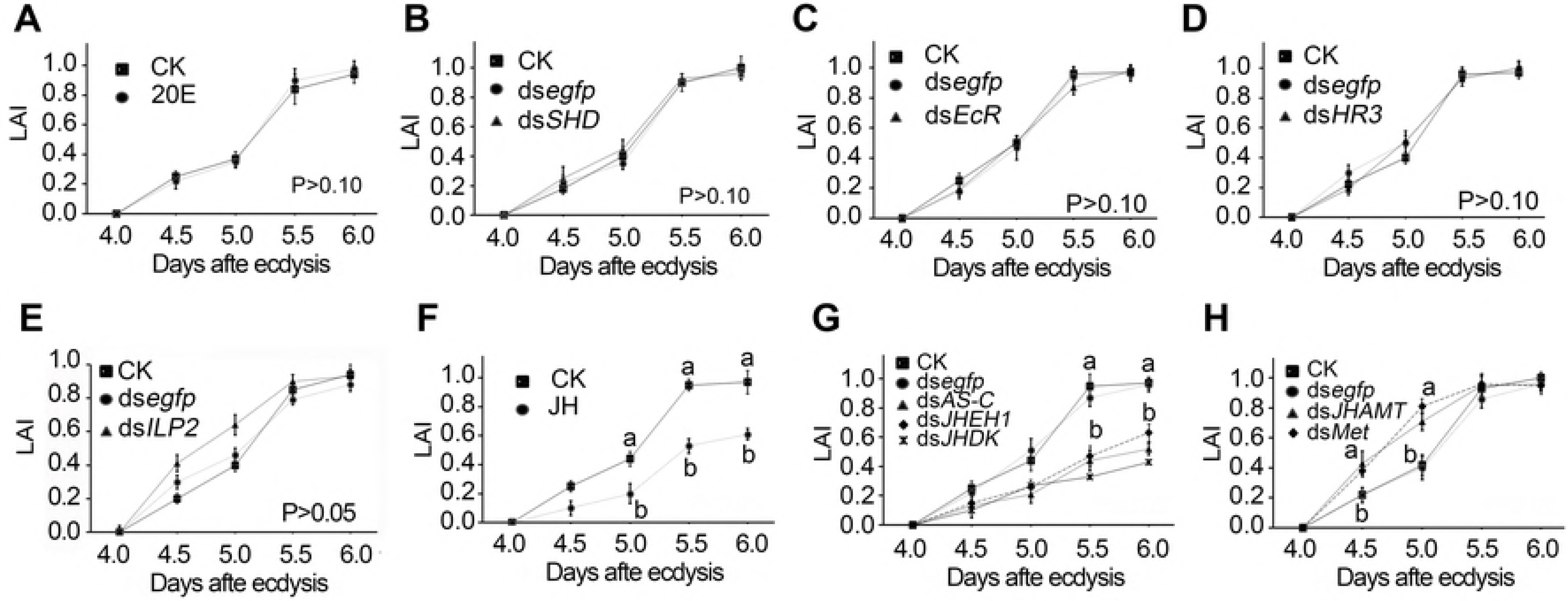
Involvement of JH signal in control of phototaxis switch. 20E pulse was prematurely generated by feeding of 20E (A); 20E signal was reduced by knockdown of an ecdysteroidogenesis gene *SHD* (B), the 20E receptor gene *EcR* (C), and a signaling gene *HR3* (D). ILP signal was inhibited by silencing of *ILP2* (E). JH titer was increased by feeding of JH (F), knockdown of *AS-C, JHEH1* and *JHDK* (G); JH pathway was repressed by silencing of *JHAMT* and *Met* (H). Larvae having ingested PBS (CK) and ds*egfp* were set as blank and negative controls. Light avoidance indexes (LAI) were calculated at each testing time point from 4.0 to 6.0 days at an interval of a half day. Significant differences between blank control (CK) and treatments at each tested time point are indicated by different letter (P < 0.05). (TIF)

Wandering behavior occurred at 4.5-4.7 days after ecdysis (S2 Table), which was earlier than the occurrence of the phototaxis switch. Moreover, ingestion of 20E accelerated the onset of wandering behavior, whereas interruption of 20E signal retarded the onset (S2 Table). The two pieces of clear evidence, different occurring time and distinct response to 20E, demonstrate that different signal cascades respectively regulate the light preference and the onset of wandering behavior.

Cessation of feeding decreased the contents of several nutrients in the larval hemolymph and thereby reduced ILP level during wandering stage [36]. Here we found that premature insulin deficiency brought about by knockdown of *ILP2* (S1D Fig) had little effect on light avoidance (Fig 2E), but delayed the onset of wandering (S2 Table).

At the wandering stage, JH titer obviously decreased [41], and the expression of JH degradation genes were activated [39]. Here we found that ingestion of JH (Fig 2F), or knockdown of an allatostatin gene (*AS-C*) [41] or either of two JH degradation genes (JH epoxide hydrolase, *JHEH1*; JH diol kinase, *JHDK*) [35, 39] to delay JH decrease significantly reduced light avoidance (Fig 2G and S1E-G Fig) and postponed the occurrence of wandering (S2 Table); whereas knockdown of a JH biosynthesis gene (JH acid methyl transferase, *JHAMT*) [34] and a JH receptor gene (methoprene-tolerant, *Met*) to prematurely reduce JH signal enhanced light avoidance (Fig 2H and S1H, I Fig) and accelerated the onset of wandering (S2 Table). It is clear that JH signal inhibits the premature switch of light preference and the early onset of wandering.

### Prothoracicotropic hormone signal acts downstream of JH

Providing elimination of JH is a prerequisite for successful PTTH signal transduction [43–45], we determined the expression levels of two PTTH signaling genes (*LdPTTH* and *LdTorso*) in the larvae whose JH signal had been disturbed. As expected, the mRNA levels of the two genes were reduced in the larval specimens whose JH signal had been enhanced (S2A-D and S3A-D Figs), and were increased in the larval samples in which JH signal had been repressed (S2E, F and S3E, F Figs).

It is suggested that PTTH promotes larval light avoidance in *L. decemlineata*. We next knocked down *PTTH* gene (Fig 3A), and verified the knockdown by determination of the mRNA level of an ecdysteroidogenesis gene (*LdPHM*) and 20E titer in the resultant larval specimens (Fig 3B). The pupation rate was decreased and the development time was lengthened in *LdPTTH* RNAi larvae; dietary supplement with 20E can rescue the two phenotypes (Fig 3C and S2 Table). Moreover, silencing of *PTTH* reduced avoidance response to light (Fig 3D), rate of larvae that had buried in soil per day (RLB) (Fig 3E) and accumulated RLB (Fig 3F). Approximately 20% of wandering larvae excavated only a slight depression at the soil surface and pupated (Figs 3G, H). Furthermore, the remaining around 80% *LdPTTH* RNAi larvae pupated at shallow layer of soil (Fig 3I); most of them did not construct pupation chambers (Fig 3J). Dietary supplement with 20E could not alleviate the reduced light avoidance response and the negative influences on pupation in *LdPTTH* depleted larvae (Figs 3D-J).

**Fig 3.**
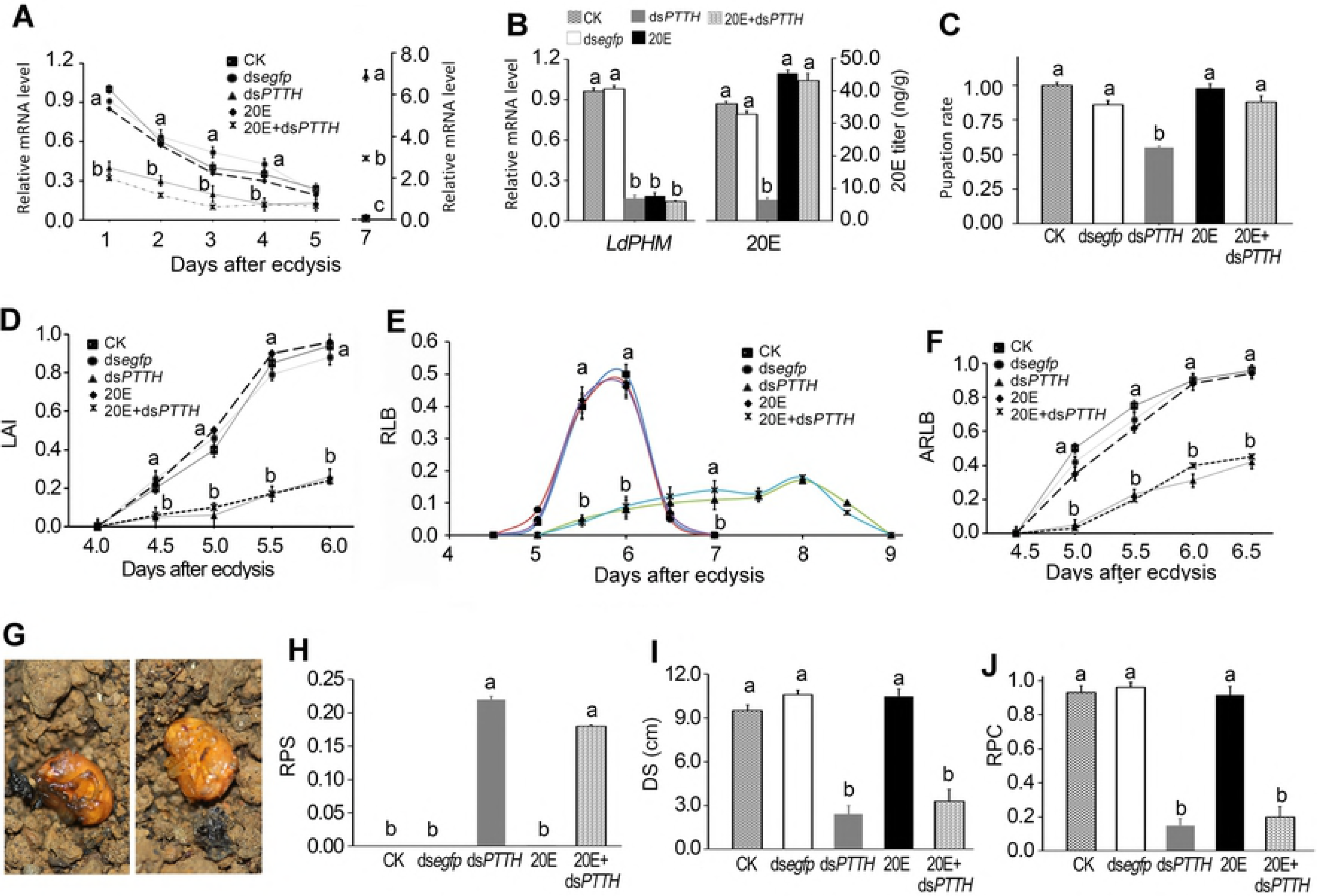
PTTH signal promotes light avoidance of the wandering larvae. PBS (CK), ds*egfp*, ds*PTTH*, 20E and ds*PTTH*+20E were dietarily introduced to the larvae. *LdPTTH* mRNA level (A), light avoidance index (LAI) (D), rate of larvae that had buried in soil per day (RLB) (E), and accumulated RBP (ARLB) (F) were determined at each testing time point demonstrating in Fig. *LdPHM* mRNA level and 20E titer (B) were tested at day 1 of the fourth-instar larvae. Pupation rate (C), rate of pupae at the soil surface (RPS) (G, H), average depths from pupation site to soil surface (DS) (I), and rate of pupae that had constructed pupation chambers (RPC) (J) were measured at the end of the experiment (P<0.05). Significant differences between blank control larvae (CK) and treatments are indicated by different letter (P<0.05). (TIF)

We repeated the bioassay by knockdown of another PTTH pathway gene, *LdTorso* [46], and obtained similar results (S3 Fig and S2 Table).

We then examined the relative mRNA levels of a JH biosynthesis gene *LdJHAMT* and two JH signaling pathway genes (*LdMet* and *LdKr-h1*), and found that these genes varied little in *LdPTTH* or *LdTorso* RNAi larvae when measured 1 and 2 days post ecdysis to fourth-instar larvae (S2G and S3Q-S Figs).

We further investigated the combination effects of PTTH and JH signaling pathways on phototaxis switch by depletion of *LdPTTH* and disturbance of JH signal (Fig 4 and S4 Fig, S2 Table). We found no additional effect on avoidance response to light (Figs 4A-D), accumulated rate of larvae in soil (Figs 4E-H) and rate of pupae on the soil (Figs 4I-L) by knockdown of *LdPTTH* and enhancement of JH signal (an addition of JH, or RNAi of *LdAS-C*), or silencing of *LdPTTH* and reduction of JH titer (RNAi of *LdJHAMT*) or JH signal (silencing of *LdMet*). Knockdown of *LdTorso* and disturbance of JH pathway mimicked the combination effects on the larval phototaxis switch (S5 Fig). These results show that JH and PTTH act on the same pathway to promote light avoidance.

**Fig 4.**
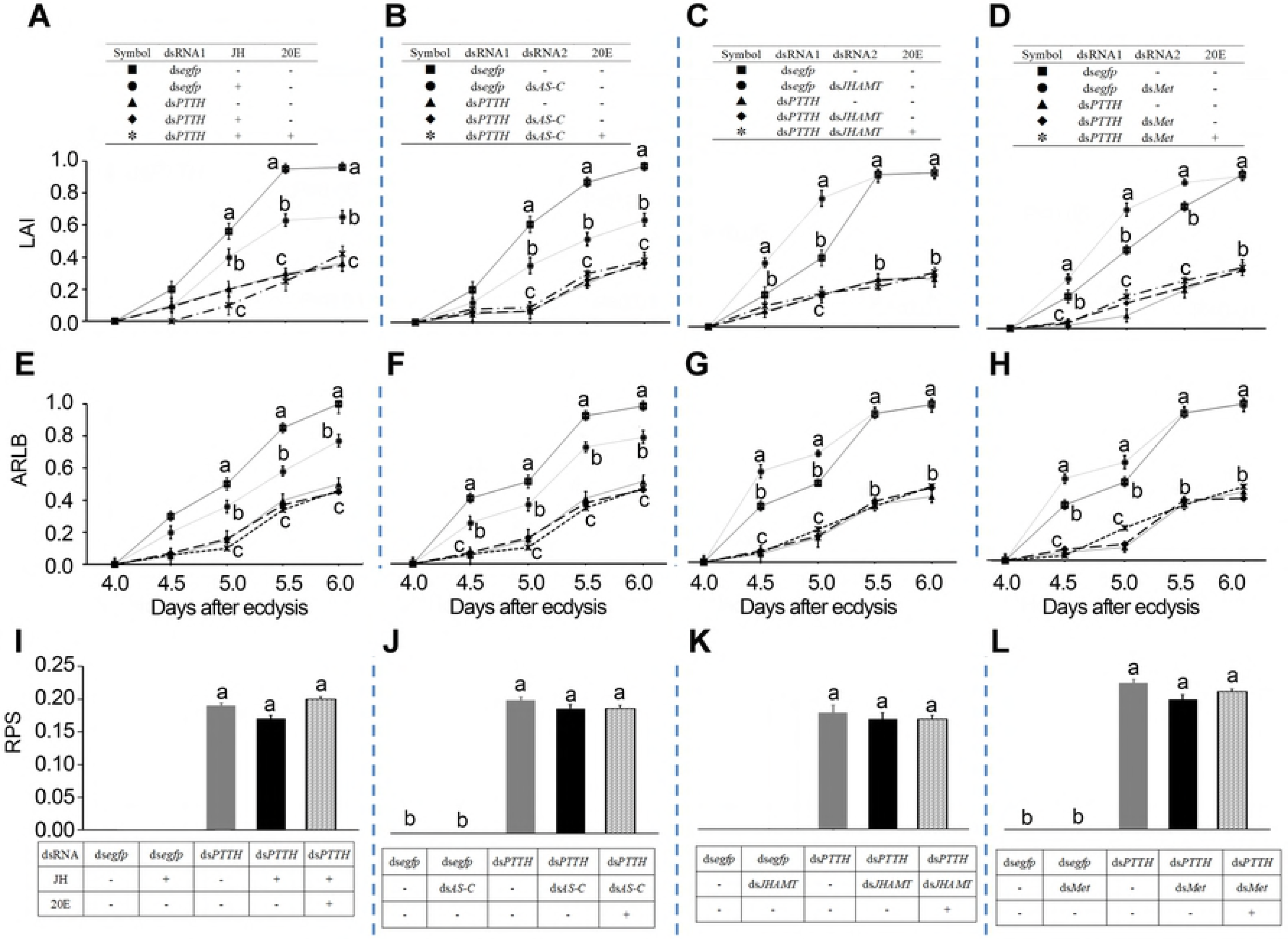
Disturbance of both PTTH and JH signals causes no additional effects on light avoidance. The larvae have fed on ds*egfp*, ds*egfp*+JH, ds*PTTH*, ds*PTTH*+JH and ds*PTTH*+JH+20E; ds*egfp*, ds*egfp+dsAS-C*, ds*PTTH*, ds*PTTH+ dsAS-C* and ds*PTTH*+ds*AS-C*+20E; ds*egfp*, ds*egfp*+ds*JHAMT*, ds*PTTH*, ds*PTTH*+ds*JHAMT* and ds*PTTH*+ ds*JHAMT*+20E; or ds*egfp*, ds*egfp*+ds*Met*, ds*PTTH*, ds*PTTH*+ds*Met* and ds*PTTH*+ ds*Met*+20E for three days. Significant differences in light avoidance index (LAI) (A-D), accumulated rate of larvae that had buried in soil (ARLB) (E-H) at each testing time point, rate of pupae on the soil (RPS) (I-L) through a two-week experiment period to those in control (ds*egfp*-fed) were indicated by different letters (P < 0.05). (TIF)

### PTTH/Torso signaling promotes light sensing

Our previous results showed that four PTTH/Torso cascade genes were highly or moderately expressed in the brain [46], suggesting that PTTH may act on neuronal cells to control light avoidance. Consistent with the suggestion, we found that the relative mRNA levels of *LdTrpA1* that encodes transient receptor potential (TRP) cation channel in *Drosophila* [47] were significantly decreased in *LdPTTH* and *LdTorso* depleted larvae. In contrast, the levels of *LdRh5* (the opsin gene involved in light avoidance behavior in *Drosophila*) [28] and *LdGr28b* (a gustatory receptor family gene that plays an opsin-like role in class IV da neurons in *Drosophila*) [48] were not affected (S6 Fig). It appears that PTTH/Torso exerts its action downstream of the photoreceptors, and upstream of TrpA1 channel activation.

Accordingly, we observed depletion of *LdTrpA1* reduced light avoidance response, and the accumulated number of larvae in soil, and increased the rate of pupae on the soil. Moreover, knockdown of both *LdTrpA1* and *LdPTTH*, or *LdTrpA1* and *LdTorso* showed no additional effects (Fig 5 and S7 Fig) (we dietarily supplemented 20E in all treatment to relieve the effect of knockdown on pupation and development time, data not shown). This provides strong evidence that PTTH/Torso signaling cascade activates a step in photo transduction between the photoreceptor molecule and the TRP channel.

**Fig 5.**
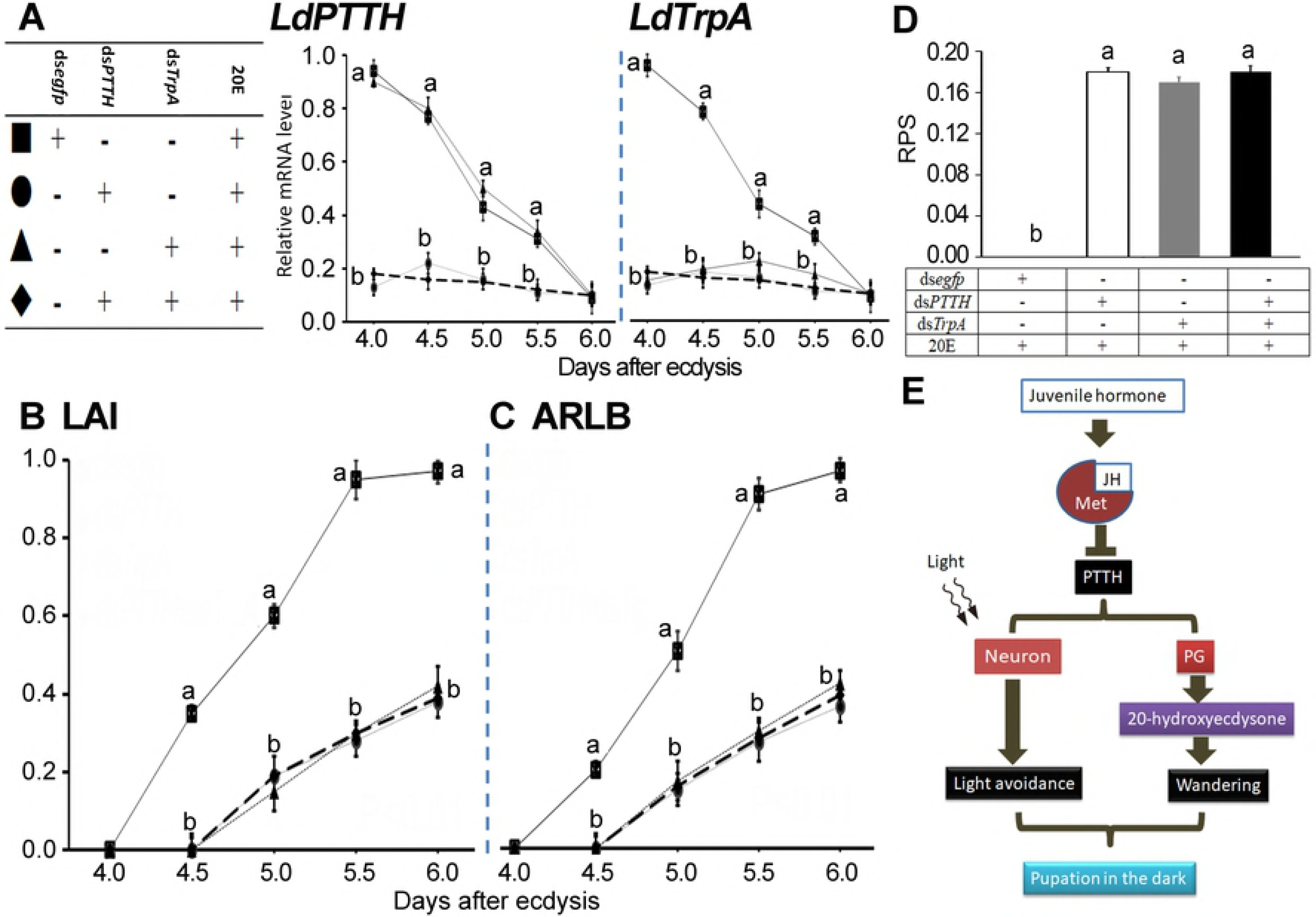
PTTH signal affects photo transduction to promote light sensing. Ds*egfp*+20E, ds*PTTH*+20E, ds*TrpA*+20E and ds*PTTH*+ds*TrpA*+20E were dietarily introduced to the larvae. Significant differences in transcript abundance of *LdPTTH* and *LdTrpA* (A), light avoidance index (LAI) (B), accumulated rate of larvae that had buried in soil (ARLB) (C) at each testing time point, and rates of pupae on the soil (RPS) (D) through a two-week experiment period to those in control (ds*egfp*+20E-fed) were indicated by different letters (P<0.05). (E) A model depicting the hormonal cascade regulating the light avoidance of the wandering larvae (see text for details).

## Discussion

Light is a vital environmental cue for living organisms [28]. In herbivorous *Leptinotarsa* larvae, we illustrate, for the first time, that the interaction of JH and PTTH signaling cascades not only regulates the onset of metamorphosis, but also promotes a phototaxis switch from photophilic at foraging stage to photophobic at middle wandering period. Therefore, JH/PTTH signaling cascade controls the decisions of when and where an herbivore undergoes metamorphosis.

### JH acts as an inhibitor for PTTH signal to suppress premature wandering and phototaxis switch behaviors

Our recent results revealed that JH signal plays an inhibitive role on PTTH production and release [43]. In this survey, we provide another three compelling pieces of evidence to support the conclusion. Firstly, we found reduced transcription levels of the two PTTH signaling genes were correlated with enhanced JH signal, and *vise verse* (S2 and S3 Figs). Secondly, JH signal concomitantly inhibited the premature switch of light preference and the early onset of wandering (Fig 2, Table 2), two behaviors can be elicited by PTTH signaling [5, 18, 21]. Lastly, no additional effects on light avoidance response and pupation were observed by simultaneous inhibition of PTTH pathway and disturbance of JH signal (Fig 4).

Insects such as *L. decemlineata, Manduca sexta, D. melanogaster* and *Trichoplusia ni*, must reach critical weigh before larval-pupal transition [43, 49–51]. When premature metamorphosis occurs below this weight, individuals tend to burden disproportionately high costs [49–52]. Therefore, it can be reasonably proposed that the presence of JH prevents premature PTTH signal, and allows insect to obtain species-specific body size, before the onset of wandering and the switch of light preference.

### PTTH-induced phototaxis switch drives *Leptinotarsa* larvae to burrow into soil

In this study, we transferred *L. decemlineata* final-instar larvae to soil at 4 days post ecdysis (the small 20E rise occurs within this day [42]). The wandering behavior occurred on 4.5-4.7 days (S2 Table). The latency between appearance of 20E and the onset of wandering is approximately 12-15 hours in *Leptinotarsa*, similar to the dormancy time in *Manduca* [22, 23]. Moreover, we found here that *Leptinotarsa* wandering larvae kept crawling at the soil surface; they did not burrow into soil (Fig 3 and S3 Fig) until the phototaxis switch had finished on day 5.5-6.0 post ecdysis (Fig 1). Therefore, PTTH-induced phototaxis switch drives *Leptinotarsa* larvae to burrow into soil.

From an ecological point of view, it seems an important evolutionary fitness for wandering *Leptinotarsa* larvae to crawl at the soil surface for an average of 24 hours before the switch of phototaxis. At the crawling period, the larvae walk a long distance away from damaged plants to complete ONS before pupation. Since herbivore-induced plant volatiles emitted by damaged plants [53, 54] attract natural enemies, maximizing distance from damaged plants prior to pupating increases the likelihood of survival. Therefore, respective regulation of the onset of wandering and the switch of phototaxis by two distinct PTTH signal branches may be a molecular evolutionary approach for insect herbivores to shift into less risky habitats for pupation.

Conversely, a portion of *Drosophila* larvae begin wandering at 108 hours after egg laying (AEL), and almost all the larvae enter wandering stage at 120 hours AEL. Comparably, some larvae start to pupariate at 108 hours AEL and almost all the larvae finish pupariation at 120 hours AEL [5]. This finding demonstrates that *Drosophila* larvae immediately form puparium even at the very beginning of the wandering period when they find an appropriate pupariation site. Consistent with the finding, the interval between PTTH release and the reinforce of light avoidance was 8 to 10 hours, while the latency time of 20E release and the occurrence of wandering is approximately 4-6 hours [55]. Considering the latency between PTTH release and ecdysone release in PGs, it is obvious that two PTTH-induced signal branches simultaneously trigger wandering behavior and reinforce light avoidance in *Drosophila* larvae.

Therefore, other cues rather than phototaxis switch decide where wandering *Drosophila* larvae pupariate. Actually, it is believed that hydrotaxis (seeking for not moisture environment) drives wandering *Drosophila* larvae to leave the food source and find a suitable pupation site [56].

### Elongated crawling period impaired pupation

It is well known that ecdysteroidogenesis in PGs are redundantly regulated by several tropic signals, such as PTTH, ILP, target of rapamycin, transforming growth factor-β/Activin and nitric oxide signals [57]. In agreement with this accepted notion, our results revealed that the wandering behavior was still activated in *PTTH* and *Torso* RNAi larvae, with a retardation of about a day (S2 Table). However, the reduced PTTH signal cannot trigger the light avoidance behavior to drive *LdPTTH* and *LdTorso* RNAi larvae to mine into soil; the beetles keep crawling for a longer period of time compared with controls. During the elongated crawling period, the enzymatic removal of dsRNA and the re-activation of transcription of *PTTH* and *Torso* finally occurred; enough mRNAs were accumulated at 7 days after ecdysis (Fig 3 and S3 Fig) and functional PTTH/Torso proteins were produced. The *PTTH* and *Torso* RNAi larvae were drove to burrow into soil, with peaked RLP values at about 8 days (Fig 3 and S3 Fig). Due to extended crawling period, *PTTH* and *Torso* RNAi beetles have less time to tunnel into soil and build their pupation chambers before the big 20E peak that elicits pupation [42]. As a result, we found in this survey that *PTTH* or *Torso* RNAi larvae pupated at the soil surface or at shallow layer of soil, with unfinished pupation chambers.

Similarly, *Manduca* larvae can construct their pupation cells when they are placed in the observation chambers during the first 20 hours of wandering. At 30 hours, larvae begin to lose their ability to complete the pupation cell. By 35-40 hours, the larvae dig only a slight depression at the soil surface before pupation [58].

Accordingly, we argue that relative constant time interval between the onset of wandering and the switch of phototaxis is crucial for an insect herbivore to accomplish ONS during the final instar to correct pupation site, a trait potentially beneficial for ecological selection. If the interval is too short, the wandering larvae have no time to choose less risky habitats for pupation. If it is too long, the wandering insects have no time to tunnel into soil and construct chambers before pupation. JH/PTTH signaling is thus at the core of a hormonal network that coordinates developmental progression and appropriate phototactic behavior to maximize insect fitness. We propose a model summarizing these findings (Fig 5E).

## Materials and methods

### Experimental model and subject details

The *L. decemlineata* beetles were kept in an insectary according to a previously described method [59], with potato foliage at the vegetative growth or young tuber stages in order to assure sufficient nutrition. At this feeding protocol, the larvae progressed through the first, second, third, and fourth instars at an approximate period of 2, 2, 2 and 4 days, respectively.

### Egg mass sampling and observation

In a 15-hectare potato field located at Urumqi city (43.82 N, 87.61 E) in the Xinjiang Uygur autonomous region of China, 100 egg masses were selected randomly and marked along a diagonal line in June 15, 2017. The development was observed and recorded at an interval of 4 hours (at night, the larvae were observed under red light) until all the larvae left the plants.

### Light/dark choice assay

The same method as previously reported [5, 27, 28] was used to test light avoidance of the larvae, with slight modifications. To synchronize the developing stage, newly-ecdysed larvae (the second through fourth instar larvae) were collected at an interval of 4 hours, and determined light avoidance at specific developing stage and different treatment, according to the experimental schedule (see Fig legend for detail). Five larvae were subjected to 20 and 30-min phototaxis assay in a Petri dish (9 cm diameter and 1.5 cm height, half of which is covered with black paper) which was illuminated from above using a white LED light at 500 lux. The larvae were placed on the center spot along the junction line between light and dark, and the larvae on which half was recorded after 20 and 30 min at a constant temperature of 25°C. Five larvae were set as a repeat, the assay repeated six times, a total of 30 larvae were determined for each instar. A steady state was reached after 20 min and we did not find any difference in the results after 20 or 30 min. The following formula for Light Avoidance (Preference) Index was used:

Light Preference Index= (number of larvae in light - number of larvae in dark) / (total number of larvae).
Light Avoidance Index= (number of larvae in dark - number of larvae in light) / (total number of larvae).

### Preparation of dsRNA

Specific primers used to clone the fragments of dsRNAs were listed in S1 Table. These dsRNAs were individually expressed using Escherichia coli HT115 (DE3) competent cells lacking RNase III following the established method [60]. Individual colonies were inoculated, and grown until cultures reached an OD600 value of 1.0. The colonies were then induced to express dsRNA by addition of isopropyl β-D-1-thiogalactopyranoside to a final concentration of 0.1 mM. The expressed dsRNA was extracted and confirmed by electrophoresis on 1% agarose gel (data not shown). Bacteria cells were centrifuged at 5000 ×g for 10 min, and resuspended in an equal original culture volume of 0.05 M phosphate buffered saline (PBS, pH 7.4). The bacterial solutions (at a dsRNA concentration of about 0.5 μg/ml) were used for experiment.

### Dietary introduction of dsRNA, 20E and JH

The same method as previously reported [60] was used to introduce dsRNA into larvae. Potato leaves were immersed with a bacterial suspension containing a dsRNA for 5 s, removed, and dried for 2 h under airflow on filter paper. The PBS-and dsegfp (enhanced green fluorescent protein)-dipped leaves were used as controls. Five treated leaves were then placed in Petri dishes (9 cm diameter and 1.5 cm height). For knockdown of *LdSHD, LdEcR* and *LdHR3*, the newly-ecdysed fourth-instar larvae were used. For other dsRNA feeding bioassays, the newly-ecdysed third-instar larvae were used. The larvae were starved for at least 4 h prior to the experiment. Then, ten larvae were transferred to each dish as a repeat. For each treatment, three repeats were set. The larvae were allowed to feed treated foliage for 3 days (replaced with freshly treated ones each day).

20-Hydroxyecdysone (20E) (Sigma-Aldrich, USA) and juvenile hormone (JH) (Sigma-Aldrich, USA) were respectively dissolved in distilled water with added surfactant (Tween 20, 1 g/L) to obtain two solutions at the concentration of 10 ng/mL. Potato leaves were dipped with 20E or JH solution. 20E supplemented leaves were provided at day 3 fourth-instar larvae, whereas JH supplemented leaves were offered at newly-ecdysed fourth-instar stage. The larvae were allowed to feed the foliage for a day.

### Real-time quantitative PCR (qRT-PCR)

Total RNA was extracted from treated and control larvae. Each sample contained 5-10 individuals and repeated three times. The RNA was extracted using SV Total RNA Isolation System Kit (Promega). Purified RNA was subjected to DNase I to remove any residual DNA according to the manufacturer’s instructions. Quantitative mRNA measurements were performed by qRT-PCR in technical triplicate, using 4 internal control genes (LdRP4, LdRP18, LdARF1 and LdARF4, see S1 Table) according to our published results [59]. An RT negative control (without reverse transcriptase) and a non-template negative control were included for each primer set to confirm the absence of genomic DNA and to check for primer-dimer or contamination in the reactions, respectively.

According to a previously described method [61], the generation of specific PCR products was confirmed by gel electrophoresis. The primer pair for each gene was tested with a 10-fold logarithmic dilution of a cDNA mixture to generate a linear standard curve (crossing point [CP] plotted vs. log of template concentration), which was used to calculate the primer pair efficiency. All primer pairs amplified a single PCR product with the expected sizes, showed a slope less than −3.0, and exhibited efficiency values ranging from 2.0 to 2.1. Data were analyzed by the 2^−ΔΔCT^ method, using the geometric mean of the four internal control genes for normalization.

### Quantitative determination of 20E and JH

20E was extracted according to a ultrasonic-assisted extraction method [40], and its titer (ng per g body weight) was analyzed by an LC tandem mass spectrometry-mass spectrometry (LC-MS/MS) system using a protocol the same as described [62].

Hemolymph was collected and JH was extracted following the methods described previously [60]. LC-MS was used to quantify JH titers (ng per ml hemolymph) [63].

### Statistical analysis

The data were given as means ± SE, and were analyzed by analyses of variance (ANOVAs) followed by the Tukey-Kramer test, using SPSS for Windows (SPSS, Chicago, IL USA), or t test. Light preference index, light avoidance index (LAI), rate of larvae that had buried in soil per day (RLP), accumulated RBP (ARLB), rate of pupae on soil (RPS), and rate of pupae that had constructed pupation chambers (RPC) were subjected to arc-sine transformation before ANOVAs.

## Acknowledgments

We are very grateful to Jiang He for assistance in insect rearing and collection. We thank other field entomologists and technicians in Urumqi for technical and other assistance.

## Supporting Information

**S1 Fig. Disturbance of 20E, ILP and JH signals**. Newly-ecdysed *Leptinotarsa* third-instar larvae had fed on ds*SHD*-, ds*EcR*-, ds*HR3*––immersed foliage for 3 days. Newly-ecdysed *Leptinotarsa* fourth-instar larvae had ingested ds*ILP2-*, ds*AS-C*-, ds*JHEH1*-, ds*JHDK*-, ds*JHAMT*- or ds*Met*-immersed foliage for 3 days. The larvae having fed PBS-or ds*egfp*-dipped foliage were set as controls. Expression levels were measured after the larvae having fed on dsRNA for three days. Significantly different mRNA levels (2^−ΔΔCt^ values±SE, the ratios of copy numbers in treated individuals relative to those in blank controls) of target and a down-stream 20E signaling gene (*LdFTZ-F1*) (A, B, C), two down-stream insulin signaling genes (*LdInR* and *Ld4EBP*) (D), or a down-stream JH signaling gene (*LdKr-h1*) (E-I), and/or 20E (A) or JH (E-H) titers were marked with different letters (P<0.05).

**S2 Fig. Transcriptional regulation of PTTH pathway by JH signaling in the final-instar larvae**. (A-F) Disturbance of JH signals influences the expression level of *LdPTTH*. JH signals were enhanced by JH ingestion (A), or knockdown of *AS-C*, *JHEH1* and *JHDK* (B-D), whereas the signals were repressed by silencing of *JHAMT* and *Met* (E, F). The expression levels of *LdPTTH* were determined. (G) Knockdown of *PTTH* did not affect the expression levels of JH signal genes. The mRNA levels of *LdJHAMT, LdMet* and *LdKr-h1* were tested at day 1 and 2 post ecdysis of the fourth-instar larvae.

**S3 Fig. Torso promotes light avoidance in the wandering larvae**. (A-F) Disturbance of JH signals influences the expression level of *LdTorso*. JH signals were enhanced by JH ingestion (A), or knockdown of *AS-C, JHEH1* and *JHDK* (B-D), whereas the signals were repressed by silencing of *JHAMT* and *Met* (E, F). The expression levels of *LdTorso* were determined.

(G-P) PTTH signal promotes light avoidance of the wandering larvae. PBS (CK), ds*egfp*, ds*Torso*, 20E and ds*Torso*+20E were dietarily introduced to the larvae. *LdTorso* mRNA level (G), light avoidance index (LAI) (J), rate of larvae that had buried in soil per day (RLP) (K), and accumulated RBP (ARLB) (L) were determined at each testing time point demonstrating in figure. *LdPHM* mRNA level and 20E titer (H) were tested at day 1 of the fourth-instar larvae. Pupation rate (I), rate of pupae at the soil surface (RPS) (M, N), average depths from pupation site to soil surface (DS) (O), and rate of pupae that had constructed pupation chambers (RPC) (P) were measured at the end of the experiment (P<0.05). Significant differences between blank control larvae (CK) and treatments are indicated by different letter (P<0.05).

(Q-S) Knockdown of *Torso* did not affect the expression levels of JH signal genes. The mRNA levels of *LdJHAMT*, *LdMet* and *LdDr-h1* were tested at day 1 and 2 post ecdysis of the fourth-instar larvae.

**S4 Fig. Disturbance of both PTTH and JH signals on gene transcription and JH titers**. The larvae have fed on ds*egfp* ds*egfp*+JH, ds*PTTH*, ds*PTTH*+JH and ds*PTTH*+JH+20E; ds*egfp* ds*egfp*+ds*AS-C*, ds*PTTH*, ds*PTTH*+ ds*AS-C* and ds*PTTH*+ds*AS-C*+20E; ds*egfp* ds*egfp*+ds*JHAMT*, ds*PTTH*, ds*PTTH*+ds*JHAMT* and ds*PTTH*+ ds*JHAMT*+20E; or ds*egfp*, ds*egfp*+ds*Met*, ds*PTTH*, ds*PTTH*+ds*Met* and ds*PTTH*+ ds*Met*+20E for three days. The expression levels of *PTTH* (A-D) and JH signal involved genes (E-H), and JH titers (I-L) were determined.

**S5 Fig. Disturbance of both *Torso* expression and JH signal impairs light avoidance**. The expression levels of *Torso* (A-D) and JH signal involved genes (E-H), and JH titers (I-L) were disturbed by allowing the larvae to feed ds*egfp*, ds*egfp*+JH, ds*Torso*, ds*Torso*+JH and ds*Torso*+JH+20E; ds*egfp*, ds*egfp*+ds*AS-C*, ds*Torso*, ds*Torso*+ ds*AS-C* and ds*Torso*+ ds*AS-C*+20E; ds*egfp*, ds*egfp*+ds*JHAMT*, ds*Torso*, ds*Torso*+ds*JHAMT* and ds*Torso*+ ds*JHAMT*+20E; or ds*egfp*, ds*egfp*+ds*Met*, ds*Torso*, ds*Torso*+ds*Met* and ds*Torso*+ ds*Met*+20E. Significant differences in light avoidance index (LAI) (M-P), accumulated rate of larvae that had buried in soil (ARLB) (Q-T) at each testing time point, rate of pupae on the soil (RPS) (U-X) through a two-week experiment period to those in control (ds*egfp*-fed) were indicated by different letters (P < 0.05).

**S6 Fig. Knockdown of LdPTTH (A-C)/LdTorso (D-F) affects the expression of light sensing genes**. Newly-ecdysed *Leptinotarsa* third-instar larvae had fed on PBS-, ds*egfp*-, ds*PTTH*-, 20E-, or ds*PTTH*+20E, or PBS-, ds*egfp*-, ds*Torso*-, 20E-, or ds*Torso*+20E-immersed foliage for 3 days. Significant differences in the mRNA levels of three light sensing genes *LdRh5, LdGr28b* and *LdTrpA1* were indicated by different letters (P<0.05). See legend in Figure 2 and S1 Fig. for further description.

**S7 Fig. Torso promotes light sensing by influence on the photo transduction**. Newly-ecdysed *Leptinotarsa* third-instar larvae had fed PBS-, ds*egfp*-, ds*Torso*-, 20E-, or ds*Torso*+20E-immersed foliage for 3 days. Significant differences in transcript abundance of *LdTorso* and *LdTrpA* (A), light avoidance index (LAI) (B), accumulated rate of larvae that had buried in soil (ARLB) (C) at each testing time point, and rates of pupae on the soil (RPS) (D) through a two-week experiment period to those in control (ds*egfp*+20E-fed) were indicated by different letters (P<0.05).

**S1 Table. Primers used in dsRNA synthesis and qRT-PCR**

**S2 Table. The period of foraging fourth-instar L. decemlineata larvae**.

## Data Accessibility

Dna sequences for this study are available in Genbank: *LdSHD* (KF044271.1), *LdPHM* (KF044261.1), *LdEcR* (AB211191.1), *LdHR3* (KP340509.1), *LdFTZ-F1* (KM091935.1), *LdILP2* (KP696397.1), *LdInR* (KT355584.1), *Ld4EBP* (KP331062.1), *LdAS-C* (KJ939426.1), *LdKr-h1* (KF137655.1), *LdJHEH1* (KP271045.1), *LdJHDK* (KP295467.1), *LdJHAMT* (KP274881.1), *LdMet* (KP147911.1), *LdPTTH* (KR152227.1), *LdTorso* (KR152228.1), *LdRas* (KR075835.1), *LdRh5* (KY368311.1), *LdGr28b* (XM_023160762.1), *LdTrpA1* (XM_023169037.1), *LdRP18* (KC190034.1), *LdRP4* (KC190033.1), *LdARF1* (KC190026.1) and *LdARF4* (XM_023168940.1).

## Author contributions

G.Q.L. and W.C.G. designed the experiments and wrote the manuscript. Q.W.M. and Q.Y.X performed the behavioral experiments. Q.W.M. and T.T.Z performed RNA interference experiments. Q.W.M. and P.D. performed quantitative RT-PCR. Q.Y.X. and K.Y.F. performed JH and 20E quantification and statistical analysis.

